# Simultaneous functional MRI of socially interacting marmosets

**DOI:** 10.1101/2021.02.08.430294

**Authors:** Kyle M. Gilbert, Justine C. Cléry, Joseph S. Gati, Yuki Hori, Alexander Mashkovtsev, Peter Zeman, Ravi S. Menon, David J. Schaeffer, Stefan Everling

## Abstract

Social cognition is a dynamic process that requires the perception and integration of a complex set of idiosyncratic features between interacting conspecifics. Here we present a method for simultaneously measuring the whole-brain activation of two socially interacting marmoset monkeys using functional magnetic resonance imaging. MRI hardware (a radiofrequency coil and peripheral devices) and image-processing pipelines were developed to assess brain responses to socialization, both on an intra-brain and inter-brain level. Notably, brain-activation maps acquired during constant interaction demonstrated neuronal synchrony between marmosets in regions of the brain responsible for processing social interaction. This method enables a wide range of possibilities for studying social function and dysfunction in a non-human primate model, including using transgenic models of neuropsychiatric disorders.

## Introduction

Social cognition is a dynamic process that requires the perception and integration of a complex set of idiosyncratic features between interacting conspecifics. Although investigations pairing unimodal stimuli with functional imaging have yielded major insights into the neural correlates of social interaction, the idiosyncrasies embedded in real social interactions—such as those communicated by reactive facial expression—are lost in these highly controlled paradigms^1^. Clever implementations of functional magnetic resonance imaging (fMRI) involving subjects interacting over a network, called hyperscanning^2-4^ has been employed where two people are simultaneously scanned in disparate scanners that are connected through an audio-video link. Hyperscanning is particularly useful for studying the unpredictability of social interactions, whereby participants’ behaviours are impacted by each other^5^. It has been noted, however, that brain activation is increased when subjects have a truly live interconnection versus watching a recorded interaction^6^. To remove the confounds of studying virtual interactions, radiofrequency (RF) coils—the hardware components responsible for receiving the MRI signal from the brain— have been developed that allow for the simultaneous imaging of two people within the same MRI scanner^7-9^. Although an elegant solution, these studies are inhibited by the limited space within the bore, which requires subjects to be in close physical contact and therefore create an unnatural social dynamic, particularly between two unrelated subjects.

Social interaction has likewise been studied in preclinical animal models, which enables the use of multi-modal and electrophysiological measurements to assess brain activation. Correlation and synchrony of neural activity during social interaction has been demonstrated in mice, bats, and non-human primates: calcium imaging of socially interacting mice has demonstrated synchrony of their neural activity predictive of social behaviour^10^; wireless electrophysiology used to record local field potentials of socially interacting bats has demonstrated correlation of neural activity over a range of timescales^11^; while neuronal ensemble recordings of non-human primates have shown inter-brain cortical synchronization during social interaction^12^. In fact, the mere presence of another monkey during the completion of a task has been demonstrated to increase brain activity in the attention frontoparietal network^13^.

Extending these animal models to represent neuropsychiatric disorders, such as schizophrenia, depression, and bipolar disorder, remains a challenging field of study^14^; however, the common marmoset monkey (*Callithrix jacchus*) is emerging as a popular animal model due to its close homology with humans in comparison to rodents^15-17^ and due to its granular dorsolateral prefrontal cortex^18^, a region of the brain that has been linked to a variety of neuropsychiatric disorders and social cognition^19, 20^. This small, New-World primate, reaches sexual maturity quickly and has a high birth rate, making it an ideal candidate for transgenic studies^21^. Marmosets can be trained to perform complex behavioural tasks while head-fixed^22^, allowing them to be used to study brain function while awake^23-27^. Recently, MRI of marmosets in the fully awake state has been performed to eliminate the confounds of anesthesia on functional activation. This technique requires specialized RF coils, such as conformal designs that clamp an individual marmoset’s head^28-30^, restraint devices with built-in RF coils^31^, and RF coils with integrated clamps to fixate an implanted chamber^32^.

Although simultaneous anatomical MRI studies have been conducted of animals (in particular, mice) for genetic studies^33, 34^, the study of socially interacting animals has yet to be investigated with fMRI—a technique which would allow a whole-brain assessment of activation in multiple animals simultaneously.

This manuscript describes a method (referred herein as the “social-coil method”) wherein hardware (including an RF coil and positioning platform) and image-processing pipelines are developed to enable simultaneous fMRI of two socially interacting marmosets on a clinical MRI scanner. The salient metrics of the coil topology are evaluated, in the context of the unique technical challenges attributed to a dual-marmoset design, to address the primary question: can the requisite image quality be achieved (in terms of signal-to-noise ratio (SNR) and limited image distortion) to map intra-brain neuronal activity and inter-brain activity of socially interacting marmosets? To demonstrate the method’s efficacy, the brain activation of two marmosets is measured during constant interaction and during a block paradigm that alters their ability to see each other. The method described in this manuscript allows for the marmoset to be adopted as an animal model to investigate whole-brain activation during social interaction in primates, enabling the study of the neural basis of social deficits in neuropsychiatric disorders using transgenic marmosets.

## Results

### Design of the radiofrequency-coil system

The radiofrequency-coil system was designed to achieve three primary goals: (1) to mitigate animal motion during functional scanning; (2) to allow marmosets to have reproducible and variant orientations within the scanner; and, most importantly, (3) to produce the requisite sensitivity for mapping brain activation on a 3T MRI scanner.

The RF coil was comprised of two disparate receive coils: each coil consisting of a marmoset restraint system with an integrated radiofrequency array. This topology was adapted from our previously published design for imaging awake-behaving marmosets on a 9.4T small-animal scanner^32^. As the limited homogeneous region of the 9.4T scanner precluded studying two marmoset brains, even in close proximity, modifications were made to allow use of our previous design on a 3T whole-body scanner and permit flexibility in the mechanical setup. The restraint system consisted of an acrylic tube equipped with neck and tail plates to constrain body motion (Fig. 1). The RF coil was affixed to the inner surface of an integrated head-fixation system, wherein the act of closing the two halves of the hinged RF coil would clamp an implanted head chamber^32, 35^ while simultaneously electrically completing the coil element circumscribing the chamber. The close-fitting nature of the receive array increases sensitivity and therefore image quality; four-point fixation of the chamber minimizes translational and rotational motion^32^.

**Fig 1.**
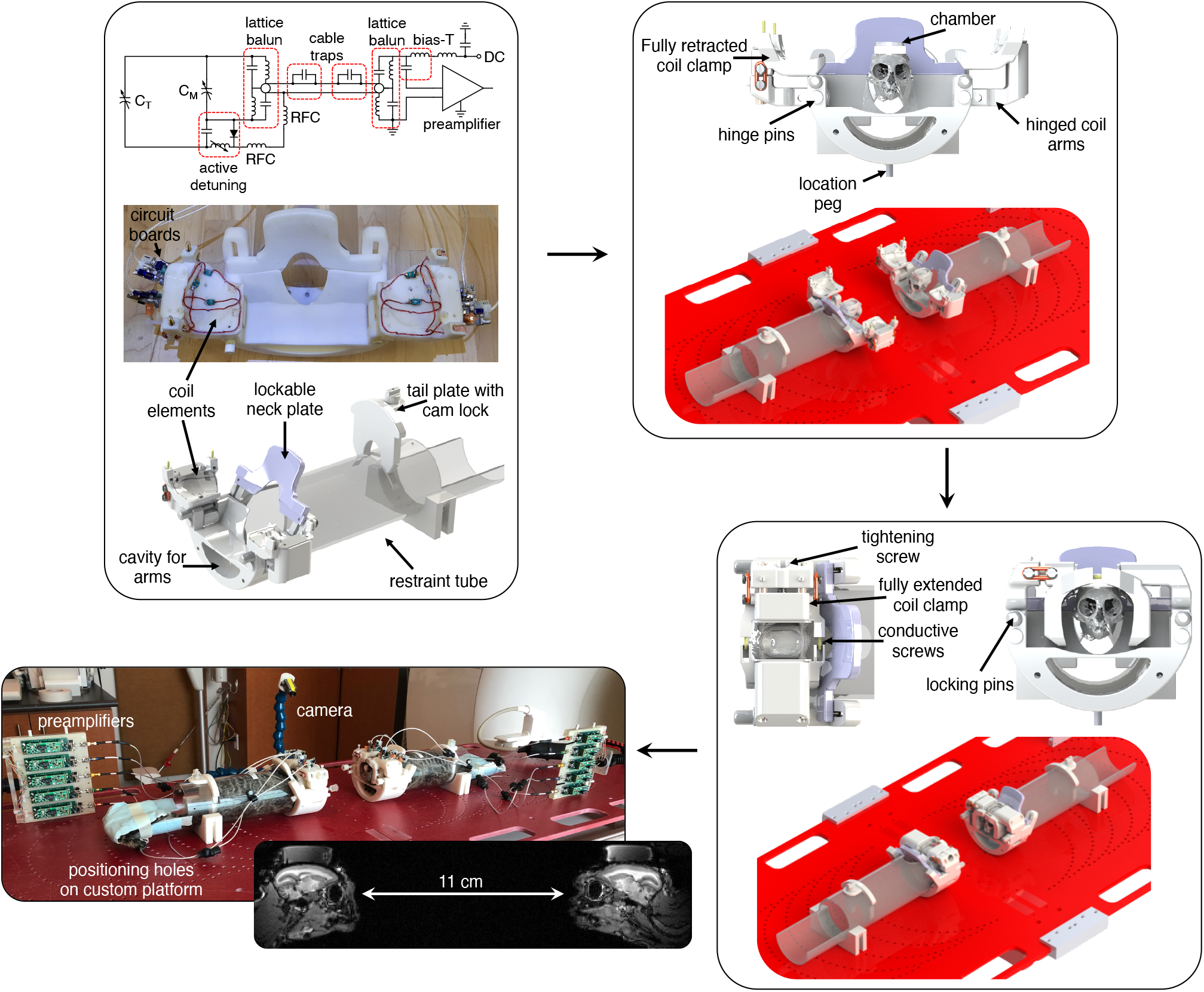
Mechanical setup of the social coil. Each marmoset is placed in a restraint device with an integrated radiofrequency coil. Initially, the arms of the restraint device are fully opened to allow the marmoset to enter a tube and be restrained by lockable neck and tail plates. Coil elements are located on the inner surface of two hinged arms; circuit boards interfacing to the coil elements are adhered to the top of the hinged arms. Once the marmosets’ bodies are restrained, they are placed on a custom platform that allows reproducible and variant positioning within the scanner. Gel is subsequently placed on the marmosets’ heads to reduce susceptibility artifacts and geometric distortion. Marmosets are then head-fixed by closing the hinged coil arms, inserting the locking pins, and fully extending the coil clamp with a tightening screw. This creates a four-point fixation of the chamber and electrically completes the coil element circumscribing the chamber by pressing conductive screws into opposing conductive pads. Preamplifier modules are repositioned independent of the coil to ensure low-noise amplifiers always maintain their optimal orientation with respect to the main magnetic field regardless of coil rotation. In this study, marmosets were placed 11-cm apart and facing each other: a distance chosen to allow natural social interaction based on the visual acuity of the marmoset. A camera was employed to monitor marmosets during scanning. C_T_: tuning capacitor; C_M_: matching capacitor; RFC: radiofrequency choke; DC: direct current.

Each receiver coil was comprised of 5 elements tuned to the Larmor frequency of protons within the 3T scanner: 123.2 MHz. Preamplifiers used in this study had the ubiquitous *B*_0_ orientation-dependence caused by the Hall effect^36^. To prevent deleterious changes to the preamplifier noise figure, and hence image SNR, long coaxial cables were used to attach preamplifiers to the coil: these enabled preamplifiers to be mounted to modules that were independent of the coil housing, allowing them to maintain the correct orientation with respect to *B*_0_ regardless of coil position. The electrical schematic of a single receive element is provided in Fig. 1.

A dedicated platform was constructed to allow reproducible and variant positioning of the two marmoset coils (Fig. 1). Two pegs underneath the coil housing could be inserted into an array of holes in the platform allowing the coil to be rotated about the *y*-axis of the scanner and translated along the *z*-axis. The confluence of the allowable translation and rotation allowed the marmosets to be set up anywhere from facing each other (allowing direct eye contact) to entirely parallel to each other (allowing both marmosets to view a common mirror or projector screen, or potentially additional animals).

### Evaluating noise characteristics of the social coil

Similarity between the performance metrics of the disparate receive coils is imperative to facilitate unbiased comparisons of brain activation between marmosets (i.e., to ensure measured differences in brain connectivity are physiological and not caused by differing coil performance). To this end, geometric decoupling, preamplifier decoupling, and active detuning were measured on the bench when coils were loaded with tissue-mimicking phantoms.

The mean and maximum coupling between receive elements was -16 dB and -12 dB, respectively (coil 1) and -22 dB and -12 dB, respectively (coil 2). The low-input-impedance preamplifiers achieved a mean decoupling per coil of -24 dB (worst-case: -20 dB), resulting in a mean and maximum *in vivo* noise correlation (Fig. 2a) of 12% and 28%, respectively (coil 1) and 13% and 29%, respectively (coil 2). The electrical and physical separation between the two coils ensures they are intrinsically well decoupled: the maximum inter-coil coupling was 2.3%. The low correlation coefficient between coils ensures that any observed synchronous brain activity between marmosets is not an artifact due to inter-coil coupling. The difference in mean noise level between coils was 9.7%, indicating similar noise characteristics were achieved through construction. Active detuning provided a minimum isolation of -29 dB between the transmit and receive coils during transmission.

**Fig 2.**
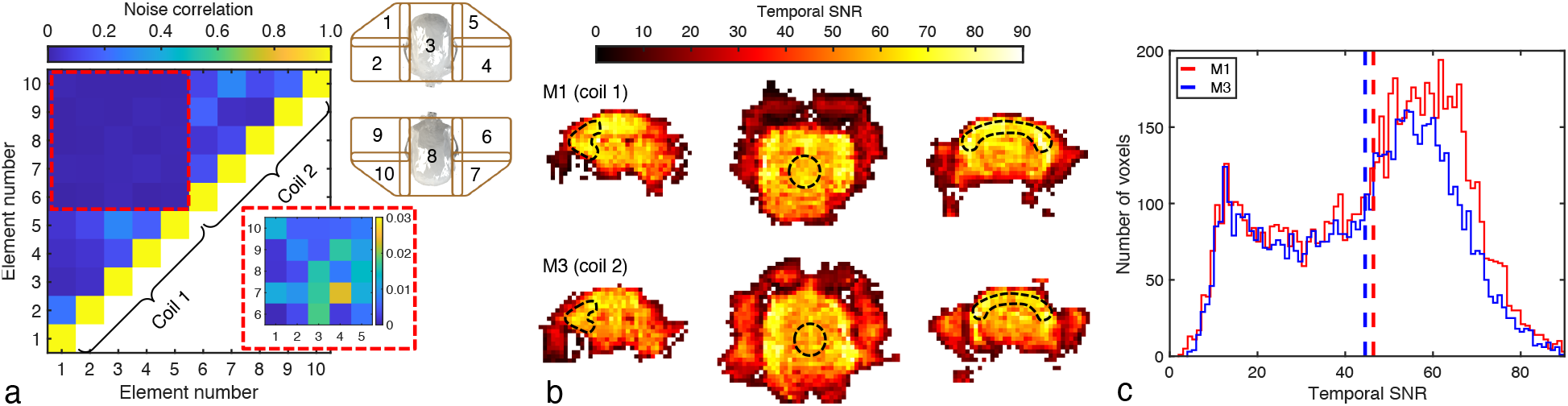
Signal and noise characteristics of the social coil. **a**, Intra- and inter-coil noise correlation of the two receive arrays corresponding to the coil-element layout and numbering depicted in the upper inset (represented in a planar view: coil 1: elements 1 – 5; coil 2: elements 6 – 10). The lower inset refers to inter-coil noise correlation, which has a maximum of 2.3%. **b**, Temporal SNR of marmoset M1 (coil 1) and marmoset M3 (coil 2) when face-to-face along the *z*-axis of the scanner. ROIs (dashed regions) in the sagittal, axial, and coronal planes show inter-brain regional differences of less than 10%, which is less than intra-brain differences (which are approximately 2 to 3-fold). Maps have been reoriented into radiological convention to facilitate comparison. **c**, Histograms of the temporal SNR of each voxel within the brain of each marmoset show similar distributions. Dashed lines represent the mean temporal SNR for each marmoset/coil combination.

### Evaluating temporal signal-to-noise ratio for functional imaging

Dedicated ultra-high-field small-animal scanners can provide high *B*_0_ fields that allow for increased SNR and image resolution^37-39^. The drawback of such systems is the typical reduction in bore size and therefore a limitation on imaging volume. Clinical scanners^40^, in contrast, can accommodate larger RF coils with more peripheral equipment and have larger imaging volumes: this can be exploited to scan multiple marmosets simultaneously, albeit at the expense of SNR (due to the lower field strength).

The temporal SNR must be high enough to accommodate a sufficient resolution to discriminate the spatial origins of the BOLD fMRI signal: this is challenging on a clinical scanner owing to the small subject size, yet large field-of-view required to accommodate two marmosets. Secondly, the two RF coils must produce consonant temporal SNR profiles to mitigate the confound of spatially varying sensitivity when quantifying synchronous brain activation.

Temporal SNR maps, derived from a 1-mm-isotropic spatial resolution echo-planar-imaging (EPI) time course, were acquired of two marmosets facing each other (Fig. 2b,c). The mean temporal SNR over the brain of each marmoset was 46.4 and 44.5, respectively—values sufficient for network mapping, as demonstrated in proceeding sections. The difference in temporal SNR between marmosets when averaged over the whole brain was 4%, whereas regional differences amounted to 9% in the frontal lobe, 7% in the centre of the brain, and 6% in the peripheral motor cortex. These small discrepancies in temporal SNR between marmosets can be attributed to both minor differences in the two RF coils as well as anatomical differences between the two marmosets.

Intra-brain heterogeneity of the temporal SNR profiles is intrinsic to all surface-coil arrays and is approximately 2 to 3-fold for the social coil. Heterogeneity in sensitivity profiles, both intra-brain and inter-brain, produce commensurate heterogeneity in the spatial sensitivity to neural correlation. To improve the accuracy of inter-brain connectivity analyses, the power of functional connectivity maps should be greater than the variability in inter-brain heterogeneity in temporal SNR, allowing for direct comparison of brain activation between interacting marmosets.

### Correcting geometric distortions on a clinical scanner

Imaging two disparate subjects within a single imaging volume reduces the efficacy of *B*_0_ shimming, with the result being a potential increase in geometric image distortion for protocols with long echo trains, such as EPI. This problem is exacerbated when trying to *B*_0_ shim over a small volume (i.e., the marmoset brain) with a whole-body gradient/shim coil; however, the lower field strength in relation to ultra-high-field small-animal scanners produces a commensurate reduction in local field inhomogeneities caused by differences in magnetic susceptibility, such as near the sinuses, the base of the skull, and regions surrounding the chamber.

No significant difference in image distortion was discerned when *B*_0_ shimming over both marmoset brains simultaneously versus over one marmoset brain at a time. Image distortion was predominantly localized to the temporal poles and the region where the chamber was affixed to the skull. This susceptibility boundary caused local off-resonance fields of up to 225 Hz, as determined with successive runs with opposing phase-encode direction^41^. This off-resonance field map was subsequently applied to correct for geometric distortion. Residual distortion was sufficiently minimal to allow accurate registration of functional images to an anatomical template^42^ (Fig. 3). The local field fidelity therefore met the threshold for reducing image distortion to a manageable level, despite the challenges of *B*_0_ shimming over two disparate and small regions-of-interest on a clinical 3T scanner.

**Fig 3.**
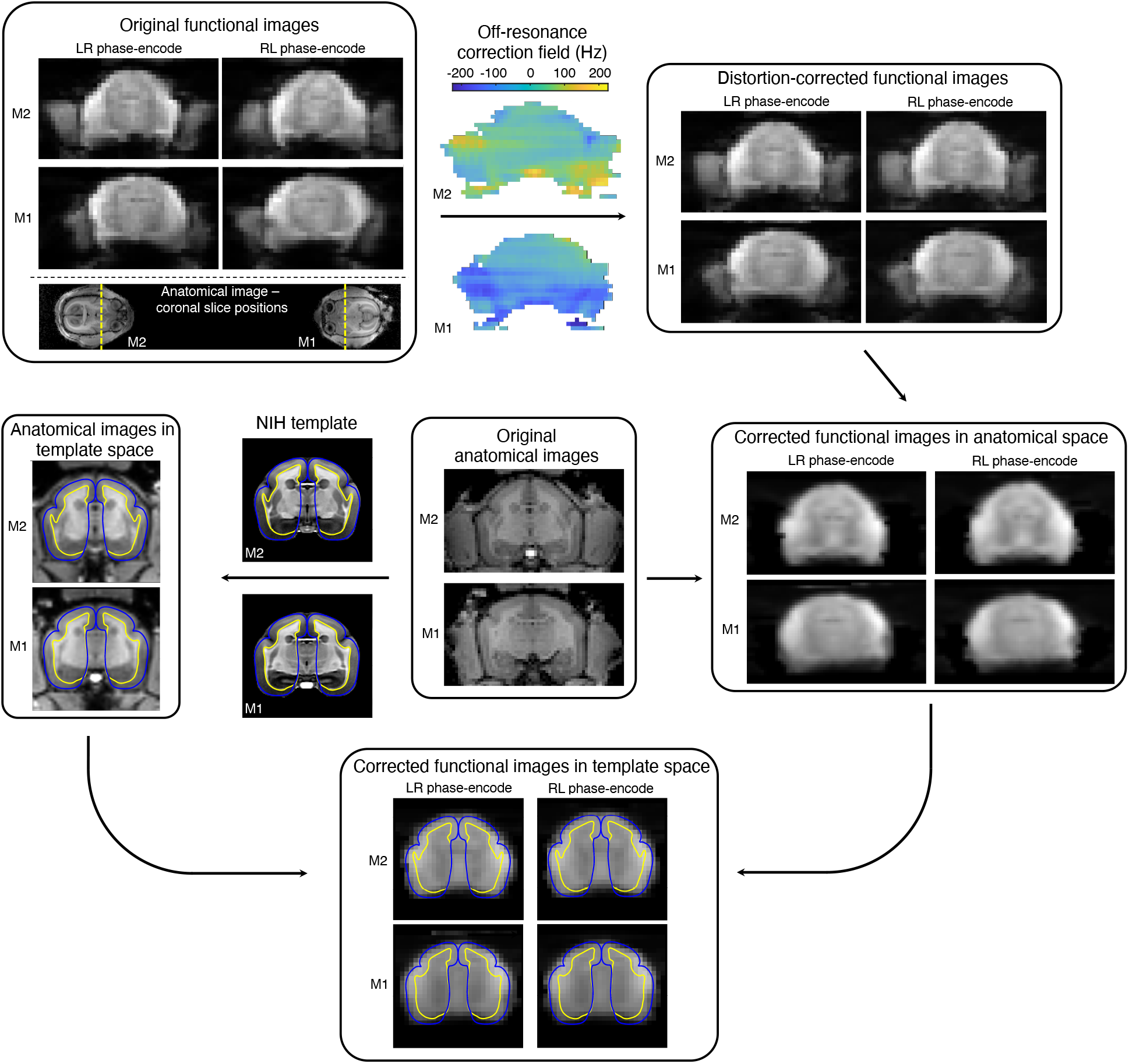
Image distortion due to local magnetic field inhomogeneities. The unique dual-marmoset setup has implications on image distortion during fMRI acquisitions: face-to-face marmosets experience opposing phase-encode directions, which creates different geometric distortion. To compensate, echo-planar images are acquired with alternating phase-encode directions. In this topology, marmoset M1, with left-right phase encode, will have similar distortion to marmoset M2 with right-left phase-encode. The most notable image distortion is found in the temporal poles and at the boundary of the chamber. Successive functional runs are then used to estimate the off-resonance correction field required for correcting image distortion. After functional images are distortion-correction and brain-extracted, they are registered to an anatomical image (i.e., anatomical space). Anatomical images, having been registered to the NIH marmoset brain atlas^61^, are used to register the functional images (in anatomical space) onto the brain atlas (template space). Registration of functional images to the marmoset brain atlas facilitates the assignment of localized brain activation to known networks. Grey- and white-matter boundaries are shown as blue and yellow lines, respectively.

### Intra-brain network mapping of constant interaction

Functional runs during constant interaction were acquired to determine whether the temporal SNR and image resolution (1-mm isotropic) would be sufficient to map intra-brain connectivity. Four runs, each 10-minutes long, were acquired with two marmosets placed face-to-face and 11-cm apart—this allowed for an uninterrupted view during the entire run. Independent component analysis was applied to each brain to determine intra-brain correlations during constant social interaction. Intra-brain connectivity maps (*z*-score maps) of three representative networks are presented in Fig. 4 (see Supplementary Fig. 1 for all derived network maps). Seven statistically significant networks were found: three somatosensory networks (SMNs), a default mode network (DMN), a primary visual network (VISp), a high-order visual network (VISh), and a salience network (SAN)— networks previously confirmed to be present in the resting marmoset^23, 32, 43^.

**Fig 4.**
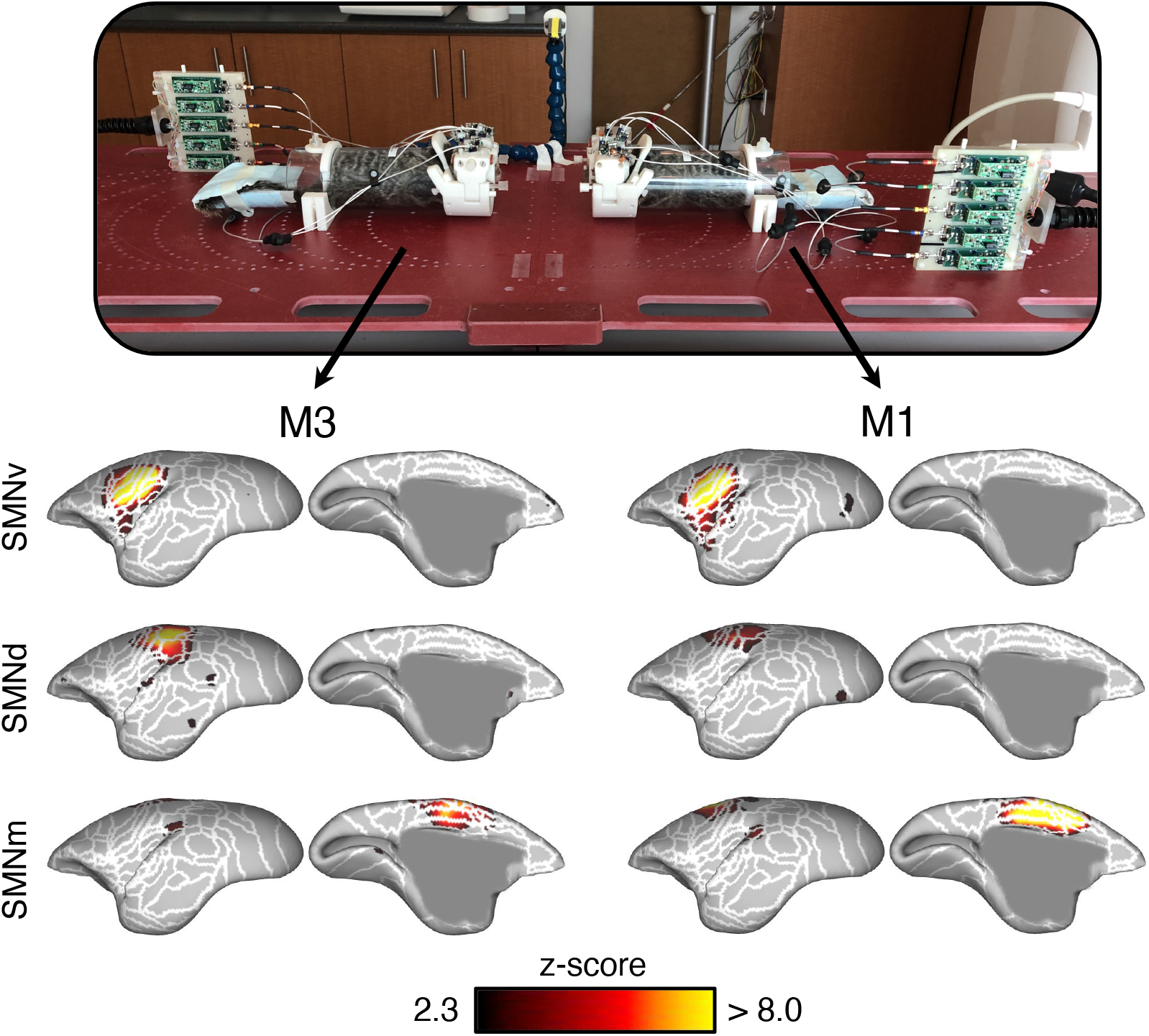
Intra-brain activation maps derived from the simultaneous scanning of two marmosets during constant social interaction. The social coil provided sufficient temporal SNR and resolution to discern seven functional networks. Three representative functional networks are displayed: the ventral somatomotor network (SMNv), dorsal SMN (SMNd), and medial SMN (SMNm). Networks are displayed as z-score maps on the cortical surface, with only the left hemisphere visible. White lines represent cytoarchitectonic borders. Supplementary Fig. 1 shows all remaining networks in both hemispheres and at the volume level.

### Intra-brain network mapping of intermittent interaction

Networks preferentially activated during socialization were evaluated by acquiring fMRI data of two marmosets during a block paradigm: the two marmosets were placed face-to-face in the dark bore of the magnet, while an LED light source intermittently illuminated the bore to allow them to see each other (Fig. 5a).

**Fig 5.**
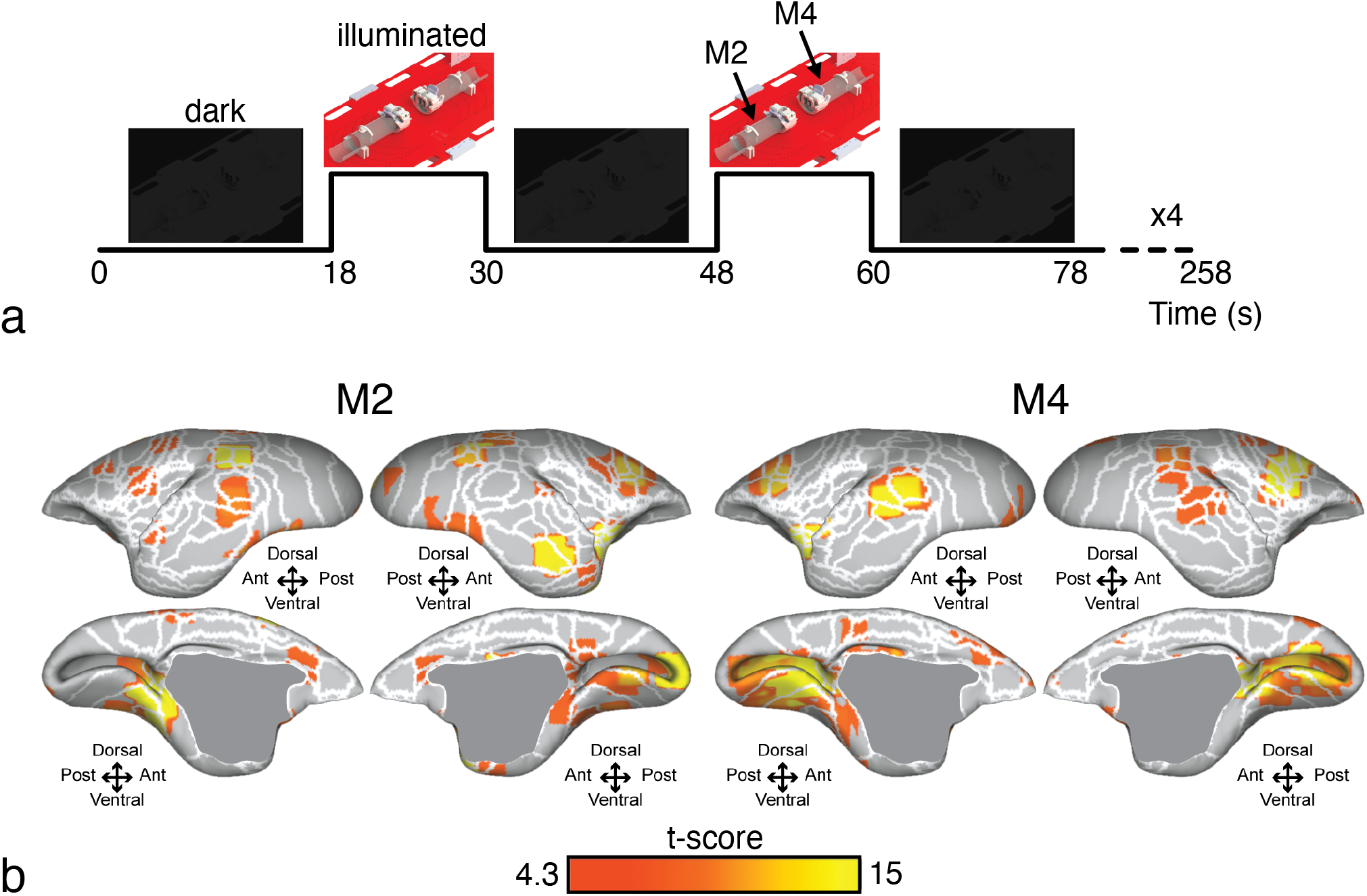
Preferentially activated brain regions during social interaction. **a**, The task-based fMRI paradigm consisted of alternating blocks wherein the scanner bore was either dark (preventing marmosets M2 and M4 from interacting) or illuminated with an LED (allowing for visual interaction).The LED was controlled through a raspberry pi that was synced to the trigger output of the scanner; the trigger occurred at the start of each image-volume acquisition. **b**, Brain-activation maps were derived by comparing the difference in activation between the two epochs of the paradigm in **a**. Activation maps are represented on the left and right fiducial surface of the marmoset brain for M2 and M4 (t-scores > 4.3, p < 0.05), with white lines representing cytoarchitectonic borders. Preferentially activated regions were found in areas of the brain associated with socialization; these regions included areas linked to visual saccades, facial recognition, and tactile stimulation.

Regions of increased brain activation during social interaction were derived by comparing the two epochs of the block design (Fig. 5b): an illuminated bore (allowing visual interaction) and a darkened bore (no interaction). Activation maps, represented on a fiducial brain surface, show significant activation occurred in multiple areas: visual (V1, V2, V3, V4, V4T), temporal (TE3, temporo-parieto-occipital association, fundus of the superior temporal, middle superior temporal, middle temporal), parietal (PG, occipito-parietal transitional area of cortex, lateral intraparietal) and frontal (dorsorostral and dorsocaudal parts of area 6, dorsal and ventral parts of area 8, caudal part of area 8, part a and b of the ventral area 6, part c of the primary motor area 4), somatosensory (1/2, 3a, 3b, S2), and cingulate (24a, 24b, 24c, 24d) cortex.

### Inter-brain synchronous connectivity

One of the unique advantages to scanning multiple awake marmosets simultaneously is the ability to assess synchronous (or conversely time-lagged and/or asynchronous) whole-brain connectivity between animals. To demonstrate the efficacy of the social-coil method, functional time-courses acquired during constant interaction were assessed for inter-brain synchrony. The *z*-score, derived from the correlation coefficient between time courses of spatially analogous voxels, showed correlated (synchronous) activity in regions A13M, A23, A24, AuCL, and V1 (Fig. 6). Of note, A24 is a region of the brain known to be involved in social-interaction processing in primates^44^. Variations in task design, marmoset pairings, and behaviour would likely evoke different regions of synchronous activity and could therefore be tailored to the neuroscientific question of interest with respect to social function and dysfunction.

**Fig 6.**
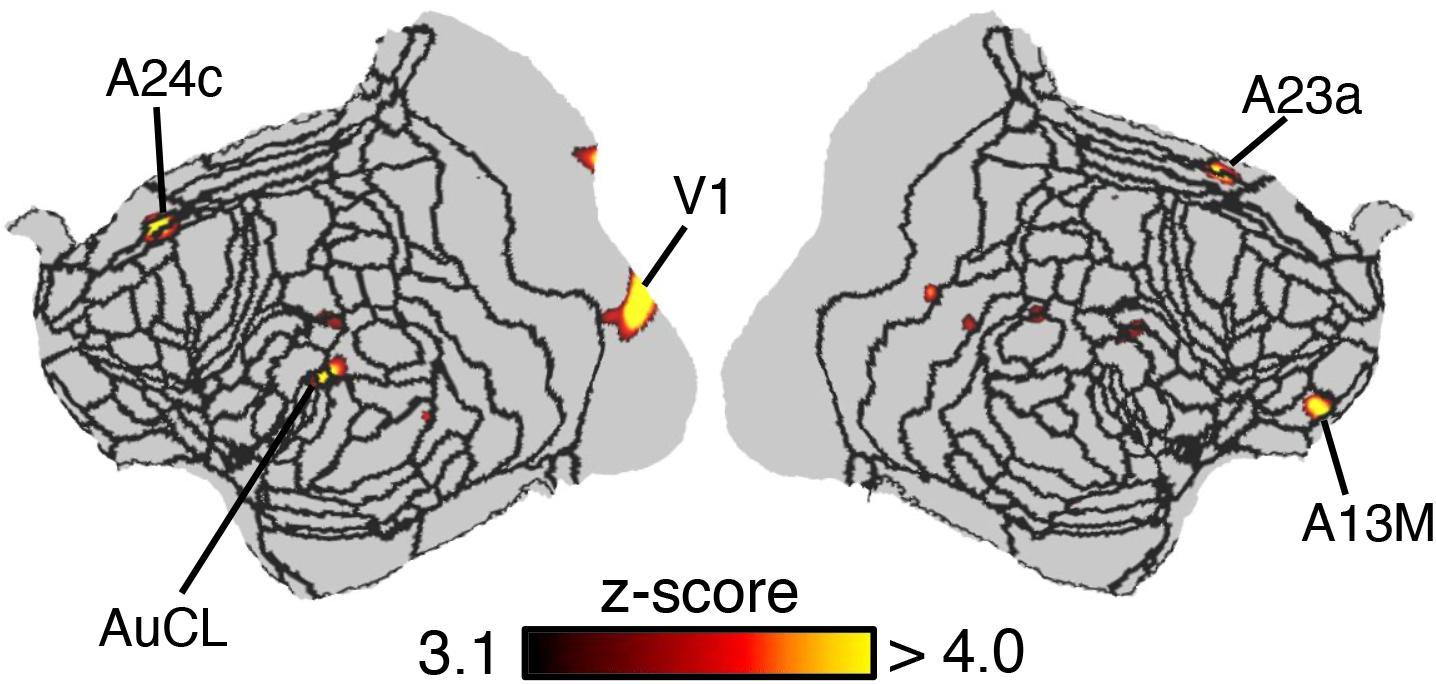
Synchronous brain activation between two marmosets during constant social interaction. Two marmosets, M1 and M3, were placed facing each other, allowing direct eye contact for the entire duration of acquiring four functional time courses. The correlation coefficient between the time courses of spatially analogous voxels in marmosets M1 and M3 was calculated and transformed to a *z*-score map—z-scores are presented on a flattened map of the left and right hemispheres, with black lines indicating cytoarchitectonic borders. Synchronous activity was found in the anterior cingulate cortex, medial prefrontal cortex, and temporal cortex—regions thought to play an important role in social interaction.

## Discussion

Investigating the brain’s reaction to socialization, using fMRI, relies on the amalgamation of dedicated hardware with image-processing pipelines. This manuscript describes a set of such tools that facilitate the simultaneous fMRI of two marmosets.

The radiofrequency coil was designed to minimize animal motion during fully awake imaging sessions, produce high sensitivity for connectivity mapping, and have the flexibility to support varying physical setups within the scanner. The confluence of these design characteristics permits the simultaneous measurement of whole-brain activation in two marmosets, enabling a variety of interactions to be investigated, from direct eye contact to the viewing of a common screen. The open-chamber design extends the flexibility of the method by allowing the integration of complimentary modalities, such as electrophysiology, to augment fMRI studies.

Operating on a 3T clinical scanner allowed for a large enough imaging region to simultaneously scan two marmosets. The caveat to using the gradient coil found in a large-bore scanner is the increased difficulty in *B*_0_ shimming over the brain regions of two marmosets; however, local susceptibility gradients in the magnetic field (and therefore geometric distortion) were sufficiently mitigated to allow accurate registration of functional images to an anatomical template with delineation between grey and white brain matter—a requisite for discriminating the spatial origins of the BOLD signal and for group analysis of multiple marmosets.

The social coil produced sufficient temporal SNR at a 1-mm-isotropic resolution to produce brain-activation maps acquired during constant and intermittent interaction. The seven functional networks obtained during constant social interaction were in agreement with resting-state networks previously observed using ultra-high-field MRI^32^. Despite the lower spatial resolution compared to typical ultra-high-field MRI studies (around 0.5-mm isotropic), the well-described small activation in the dorsolateral frontal area in DMN^45, 46^ was detected. Nonetheless, increasing the number of coil elements would facilitate higher acceleration rates, resulting in reduced geometric image distortions and permitting higher resolution functional data—an achievable goal due to the high temporal SNR.

Brain-activation maps generated by the intermittent-interaction paradigm demonstrated regions of the brain preferentially activated during social interaction. These included networks previously identified in marmosets performing tasks^25, 27, 47^, but most notably, activation was found in regions associated with social cognition. Activation linked to the visuo-saccadic network of marmosets^27^ (frontal, parietal, temporal, and visual regions of the brain) was observed, as marmosets in visual contact will perform saccades to analyze each other’s face^22^. Most likely due to the same mechanism, regions typically activated during facial scanning^19, 25^ (i.e., face-patch regions in the temporal cortex: AD, MD, PD, and PV) were also preferentially activated. Furthermore, activation was observed in areas of the brain that activate in response to light tactile stimulations to the face^47^ (lower part of somatosensory areas 3a, 3b, 1/2, as well as in the rostral part of the somatosensory area S2). Despite the marmosets’ inability to make physical contact during scanning, their close proximity may have elicited an anticipation response to potential interaction or contact. The somatosensory cortex has also been linked to the mirror system neuron^48-51^, theory of mind, emotions, and empathy in humans and non-human primates. Taken together, these results indicate that the social-coil method has the sensitivity to discriminate the brain’s response to social interaction. Importantly, this can be performed at the whole-brain level.

Most notably, the social-coil method was capable of revealing synchronous neuronal activity between two marmosets. Synchronicity was found in regions thought to play an important role in social interaction (the anterior cingulate cortex (ACC), medial prefrontal cortex (mPFC), and temporal regions). For example, these brain regions in macaque monkeys have shown increased activity when they watch movies showing monkeys interacting versus acting independently^44^. In humans, mutual interaction during eye contact has been shown to be mediated by the limbic mirror system, including the ACC and anterior insular cortex (AIC)^52^. Furthermore, inter-brain synchrony has been found in regions including the ACC and right AIC when playing a modified interactive ultimatum game^53^. Taken together, correlated activity in area 24 (a part of the anterior and middle cingulate cortex) and the temporal region infers a synchronization of regional brain activity due to the live interaction (socialization) between the marmosets. The ability for the social-coil method to detect synchronous activity allows for the study of increasingly sophisticated animal models and social interactions.

The method described in this study enables the measurement and assessment of synchronous neuronal activation, across the whole brain, between interacting marmosets. This allows the study of social function and dysfunction in a non-human primate model, including the use of transgenic models of neuropsychiatric disorders. The first demonstration of simultaneous functional imaging of two marmosets within the same scanner removed the confounds of remote hyperscanning. The modular coil design can now be expanded to allow for the simultaneous scanning of larger cohorts, enabling a broad range of social groups and interactions to be investigated.

## Methods

### Social-coil hardware topology

The social coil was comprised of a marmoset restraint system with an integrated radiofrequency coil (Fig. 1). This system was adapted from our previously published design for imaging awake-behaving marmosets on a 9.4T small-animal scanner^32^.

The body-restraint consisted of a 7-cm-diameter acrylic tube (product number: 8532K23; McMaster-Carr, Aurora, Ohio, USA) with neck and tail plates. Marmosets would enter the tube, be restrained by an adjustable neck plate with thumb screws and a tail plate with cam lock: the neck plate and receive coil were angled to accommodate the marmoset’s body. The marmoset would then lie in the sphinx position with the coil arms fully opened. Once acclimatized to being in the body-restraint, marmosets would be head-fixed by closing the coil arms and locking them in place by inserting hinge pins. The retractable clamp is then tightened, by turning a screw, to secure the fixation pins into the chamber. The head-fixation assembly was 3D-printed using stereolithography with a photopolymer resin (White Resin V4; Form 2, Formlabs, Somerville, Massachusetts, USA).

### Design and fabrication of the integrated receive array

The receive array included five loops integrated into the interior of the coil former to allow close proximity to the brain (and therefore higher SNR). One element circumscribed the head chamber, while two elements were arranged along the anterior-posterior direction on either side of the head: this allowed for a two-fold acceleration rate in the anterior-posterior or left-right directions during parallel imaging. Coil elements were geometrically overlapped to reduce inter-element coil coupling^54^. Each element was constructed from 22-AWG insulated copper wire and included three or four distributed capacitors to reduce conservative electric fields: one variable capacitor for matching (Sprague-Goodman Electronics, SGC3 series), one fixed capacitor for the active-detuning circuit, and one variable capacitor for tuning (an additional surface-mount capacitor was incorporated into the large element circumscribing the chamber). Each element was tuned to 123.2 MHz and matched to 200 + j50 Ω (i.e., the optimal noise impedance of the preamplifier) when loaded with a 2.5-cm-diameter spherical phantom filled with 50-mM sodium chloride. Polyurethane foam (product number: 86375K162; McMaster-Carr, Aurora, Ohio, USA), 1.6-mm-thick, was adhered to the inner surface of the coil as a spacer to prevent conservative electric fields from coupling to the marmoset (which would result in a shift in the resonant frequency and a reduction in SNR).

The element circumscribing the chamber was split into two halves. When tightening the retractable clamp to secure the chamber, two conducting posts (6-32 brass screws) on one arm pushed into two conductive pads on the other arm to create electrical continuity. The conductive pads were created by wrapping copper braid in front of ethylene-vinyl-acetate (EVA) foam and soldering the braid to one half of the coil element. The flexibility of the foam created solid connections between the conducting posts and associated pads to decrease coil noise and prevent spiking artifacts.

Circuit boards were mounted on the outside of the former and included a matching capacitor, an active-detuning circuit, and a lattice balun. The low-input-impedance preamplifiers (Stark Contrast, Erlangen, Germany) had a *B*_0_ orientation-dependence due to the Hall effect. To prevent a reduction in preamplifier noise figure, 61-cm-long RG178 coaxial cables (i.e., a half-wavelength electrical length) attached the coil to the preamplifiers through non-magnetic MCX connectors. Preamplifiers were then mounted to modules that were independent of the coil housing, yet movable themselves, and consistently orientated with the low-noise amplifier parallel to *B*_0_. The longer cables required additional cable traps to suppress common-mode currents: in addition to the lattice baluns placed at the input of the coil, a lattice balun was located at the input of the preamplifier (to ensure a symmetric input to the preamplifier) and two choke baluns were placed along the coaxial cable. The second lattice balun had the opposite orientation to the lattice balun at the coil input to counteract any impedance transformation. The baluns were sufficient at preventing changes in the tune and match of coil elements with varying cable position. The half-lambda cables transformed the low input impedance of the preamplifier to a parallel-resonant inductance across the matching capacitor, creating a high-impedance circuit to reduce inter-element coupling (i.e., preamplifier decoupling^54^). A multi-conductor cable connected preamplifiers to the system socket; this cable had a split sleeve balun to reduce common-mode coupling to the transmit coil.

### Design and fabrication of a custom positioning platform

A dedicated platform was constructed to allow reproducible positioning of the two marmoset coils (Fig. 1). The platform was comprised of 0.95-cm-thick garolite with legs that slotted onto the scanner bed. Slots were machined on the side of the platform to allow for ease of handling.

Two pegs underneath the coil housing could be inserted into an array of holes in the platform: each coil could be rotated about the *y*-axis of the scanner by up to 180°, in 5° increments; the *z*-position of each coil could be varied in 5-cm increments to allow approximately 1 – 91 cm of space between marmosets. Marmosets could be oriented to allow face-to-face interaction with direct eye contact to being entirely parallel to each other to view a common mirror or projector screen. Since the sensitivity to transverse magnetization decreases with increased angle to *B*_0_, coils should be placed at conjugate angles with respect to *B*_0_ to avoid introducing an SNR disparity between the two coils.

### Bench evaluation of the social coil

Standard single- and double-probe techniques^55^ were employed to measure geometric decoupling, preamplifier decoupling, and active detuning on the bench using a network analyzer. Geometric decoupling was measured as the transmission coefficient between preamplifier ports. Preamplifier decoupling and active detuning were measured as the difference between the tuned state (i.e., without a preamplifier present or detuning bias) and when the coil had a preamplifier present and was tuned (preamplifier decoupling) or detuned (active detuning). Coils were loaded with 2.5-cm-diameter spherical phantoms filled with a 50-mM sodium-chloride solution to approximate physiological conductivity.

### The marmoset animal model

Experimental procedures were in accordance with the Canadian Council of Animal Care policy and a protocol (#2017-114) approved by the Animal Care Committee of the University of Western Ontario Council on Animal Care. Imaging was performed on four common marmosets: 3-year-old males weighing 310 g (M1), 400 g (M2), and 340 g (M3), and a 2.5-year-old female weighing 365 g (M4).

Marmosets underwent an aseptic surgical procedure^35^ to implant a head chamber while the animal was placed in a stereotactic frame (Narishige, Model SR-6C-HT). Adhesive resin (All-bond Universal, Bisco, Schaumburg, Illinois, USA) was applied using a microbrush, air dried, and cured with an ultraviolet dental curing light (King Dental), after which a two-component dental cement (C & B Cement, Bisco, Schaumburg, Illinois, USA) was applied to the skull and the bottom of the chamber. The chamber was then lowered onto the skull with a stereotactic manipulator to ensure accurate placement. A 3D printed cap was attached to the chamber with set screws; this cap was removed before entering the magnet room.

All animals were acclimatized to the restraint system and coil prior to imaging^32, 56^. This procedure required three weeks and included being placed in the restraint system for increasingly long durations and being played MRI sounds at a loud volume.

Directly prior to scanning, marmosets were placed in the restraint system, but not head fixed, in a preparation room adjacent to the magnet room. Marmosets were head-fixed once on the scanner bed to minimize risk during transfer. During imaging, an MRI-compatible camera was used by a veterinary technician to monitor marmosets for stress and to check as to whether the animals were awake.

### MRI scanner hardware

All imaging was performed at the Centre for Functional and Metabolic Mapping at The University of Western Ontario. MRI data collection was performed on a human, whole-body Siemens Prisma Fit scanner (Siemens Healthineers AG, Erlangen, Germany) operating with a 3-T main magnetic field. The system is equipped with 64 receiver channels, of which 10 were utilized: two plugs (one per coil) were interfaced to Tim coil interface adaptors to allow operation on the Siemens Prisma hardware platform. The SC72 gradient coil allowed for a maximum gradient strength of 80 mT/m and a maximum slew rate: 200 T/m/s.

### Measuring the temporal signal-to-noise ratio

A gradient-recalled-echo, noise-only acquisition (i.e., without RF transmission) was acquired to calculate the noise level and noise correlation matrix of and between the two disparate coils (matrix size: 384 Ω 156, FOV: 256 Ω 104 mm, TE/TR: 4.6/10 ms, BW: 260 Hz/pixel).

Temporal SNR maps were calculated from a single-shot, EPI time series with two marmosets facing each other: FOV: 220 Ω 78 mm, acquisition matrix: 220 Ω 78, number of slices: 25, slice thickness: 1 mm, TE/TR: 30/1,500 ms, BW: 1,265 Hz/pixel, flip angle: 70°, volumes: 400, acceleration rate: 2 (right-left), reference lines: 24, partial Fourier encoding: 6/8. Temporal SNR maps were calculated by measuring the ratio of the mean signal of each pixel through the de-trended time course to the standard deviation of that pixel through the time course. Temporal SNR calculations were performed in Matlab (The MathWorks, Natick, MA).

ImageJ was used to calculate the mean temporal SNR in regions of interest located in the frontal lobe, centre of the brain, and peripheral motor cortex. The mean temporal SNR was also calculated over the entire brain region (i.e., excluding the temporal muscles, eyes, etc.).

### Assessing geometric distortion

Head chambers often produce local image-intensity dropouts due to differences in the magnetic susceptibility between the chamber, air, and tissue. Susceptibility-induced distortion attributed to the chamber was partially abated by applying a water-based gel (MUKO SM321N; Canadian Custom Packaging Company, Toronto, Canada) to the brow ridge and inside the chamber, thus reducing susceptibility differences close to the brain.

Marmosets M1 and M2 were placed facing each other 11-cm apart—a distance chosen based on the visual acuity of marmosets^57^. *B*_0_ shimming was performed over a volume large enough to encompass both marmoset brains. Single-shot, multiband^58^ EPI functional runs were acquired of both marmosets, simultaneously (orientation: axial; FOV: 220 Ω 78 mm, acquisition matrix: 220 Ω 78, number of slices: 25, slice thickness: 1 mm, TE/TR: 30/1,500 ms, BW: 1,265 Hz/pixel, flip angle: 70°, acceleration rate: 2, reference lines: 24, partial Fourier encoding: 6/8). Accelerated images were reconstructed with generalized autocalibrating partially parallel acquisition (GRAPPA)^59^.

Since marmosets have opposite orientations within the scanner, their respective image distortions, caused by local field inhomogeneities, have opposing anatomical directions. To correct for this difference, successive functional datasets were acquired with opposite phase-encode direction (left-right versus right-left).

From these successive runs, the susceptibility-induced off-resonance field was estimated^41^ and used to correct for distortion (FSL^60^; topup). A magnetization-prepared, rapid gradient echo (MPRAGE) image was acquired as an anatomical reference (FOV: 220 Ω 68; matrix: 352 Ω 110, number of slices: 64, slice thickness: 0.63, TE/TI/TR: 4.2/900/2300 ms, BW: 200 Hz/pixel, flip angle: 9°, number of averages: 2). Functional images were registered to the NIH marmoset brain atlas^61^ as described in the proceeding section.

### Intra-brain network mapping of constant interaction

Functional time courses (echo-planar images) were acquired during constant interaction (i.e., while marmosets could make direct, uninterrupted eye contact). Four runs, with 400 volumes each, were acquired with marmoset M1 and M3 facing each other and placed 11-cm apart. Pulse-sequence parameters were identical to those described in the previous section, with alternating phase-encode direction along the left-right axis (i.e., left-to-right versus right-to-left).

Prior to functional runs, *B*_0_ shimming was performed over the imaging volume encompassing both marmosets. An anatomical reference scan (the MPRAGE sequence) was acquired of the multiple marmosets for image registration (pulse-sequence parameters were identical to those described in the previous section). Functional data was pre-processed and analyzed using an adaptation of our previously published method^32^, as described below.

Anatomical images were split into separate datasets for each marmoset. These datasets were then reoriented to radiological convention to compensate for the different orientation of each marmoset within the scanner. The process of removing the skull from the anatomical image was conducted in three stages: (1) the olfactory bulb was manually removed, as it was not included in the template image; (2) the delineation of the brain-skull boundary was approximated (FSL; BET; radius: 25-30 mm; fractional intensity threshold: 0.3); and (3) the *T*_1_-weighted brain template^61^ was linearly and nonlinearly registered to the skull-stripped anatomical image (FSL; FLIRT and FNIRT) to refine the brain-skull boundary, create a mask, and create an atlas-to-anatomical transformation matrix.

Functional images were split to have one distinct dataset per marmoset. Data from each marmoset was preprocessed separately using FSL^60^. As with anatomical images, functional images were reoriented to the standard radiological convention. Functional images were then corrected for motion (FSL; FLIRT) and geometric distortion (FSL; topup) and manually brain extracted (FSL; fslview). An average functional image was calculated for each run and an anatomical-to-functional transformation matrix was calculated (FSL; FLIRT and FNIRT).

Functional images (still in native space) were finally normalized to the template using the inverse of the transformation matrices (functional-to-anatomical and anatomical-to-atlas)—this facilitated the assignment of brain activation to known brain regions. This was followed by spatial smoothing (1.5-mm full-width at half-maximum Gaussian kernel) and temporal filtering (0.01 to 0.1 Hz) (FSL; fslmaths).

Principal component analysis (PCA) was applied to remove the unstructured noise from each animal’s time course (FSL; MELODIC), followed by independent component analysis (ICA) with 100 dimensions. The resultant components were classified as signal or noise based on the criteria as shown in previous reports^62^. Noise components were regressed from each fMRI time course (FSL; fsl_regfilt). Group ICA, with 20 dimensions, was subsequently performed on each marmoset’s data to detect the neural components.

### Intra-brain network mapping of intermittent interaction

Regions of the brain preferentially activated during social interaction were deduced by measuring the difference in marmosets’ brain activation between periods with and without the ability to view one another. Two marmosets (M2 and M4) were placed face-to-face, at a distance of 11 cm, within the blacked-out bore of the MRI scanner. In a block paradigm, an LED was illuminated to alternate the animals’ ability to see each other. Each run was comprised of 17 alternating blocks: 18 s in the dark condition (i.e., no interaction) followed by 12 s in the illuminated condition (i.e., freely viewing each other). The LED was controlled by a Raspberry-Pi (Raspberry-Pi 3, Model B), through a Python script, which was synced to the trigger output of the scanner that was sent at the beginning of each volume acquisition.

Functional (EPI) and anatomical (MPRAGE) data were acquired with the same acquisition parameters as described for constant-interaction runs, albeit with 172 volumes per run. Two runs were acquired in one session: one with left-right acceleration and one with right-left acceleration.

Task-based data was pre-processed using primarily the same pipeline as described in the previous section, with small differences pertaining to the processing of functional images. After image reorientation, motion correction, and distortion correction, functional images (in native space) were despiked (AFNI; 3dDespike) and volume registered to the middle volume of each time series (AFNI; 3dvolreg). Images were smoothed by a 2.0-mm full-width at half-maximum Gaussian kernel to reduce noise (AFNI; 3dmerge).

Task timing was convolved with the hemodynamic response function (AFNI; BLOCK convolution), and for each run a regressor was generated for each condition to be used in a regression analysis (AFNI; 3dDeconvolve). Both conditions were entered into the same model, along with fifth-order-polynomial detrending regressors, bandpass regressors, and the motion parameters derived from the volume registration. The resultant regression-coefficient maps were then registered to the NIH template space^61^ using the transformation matrices described in the previous section. Marmoset t-value maps were then compared at the individual level using a t-test (AFNI; 3dttest++); resultant t-values were displayed on fiducial maps using Connectome Workbench^63^ in conjunction with the NIH marmoset brain template^61^.

To ensure the switching of the LED did not alter the received signal, scans were acquired without marmosets (i.e., noise-only images) using the identical paradigm—no difference in noise level was observed between states of the LED, nor were noise spikes observed during the switching of the LED.

### Inter-brain synchronous connectivity

Inter-brain synchronous connectivity was deduced by calculating the voxel-wise correlation coefficient between the simultaneously acquired functional time series of two marmosets during constant visual interaction. Constant-interaction time-course data was preprocessed as described previously and voxel intensities were subsequently normalized to have a mean of zero and a standard deviation of 1. (This latter step mitigated bias due to inter-run differences in the mean and standard deviation of voxel intensities.) Time courses of the four independent runs were temporally concatenated on a voxel-wise basis. The correlation coefficient was then calculated between spatially analogous voxels in the disparate marmoset brains. Correlation-coefficient maps were Fisher-Z transformed and masked to a threshold of 3.1, corresponding to *p* = 0.01 when corrected for multiple comparisons using Gaussian Random Field theory. Calculations were performed in Matlab (The MathWorks, Natick, MA).

### Functional data acquisitions

EPI datasets acquired in this study employed imaging volumes large enough to encompass both marmosets, which requires the acquisition of non-encoded space and therefore increased noise. This, however, is a limitation imposed by the pulse sequence and is a tractable problem. Multi-animal functional MRI would lend itself well to simultaneous multi-slice imaging techniques^64^. With the modification to allow for two stacks of slices, a slice acceleration factor of two-fold would allow for simultaneous acquisition of a single slice in each marmoset acquired from disparate, decoupled RF coils. Consequently, the distance between marmosets could be increased without a proportional increase in imaging volume, and commensurately the SNR should improve.

## Acknowledgements

The authors would like to thank Cheryl Vander Tuin and Hannah Pettypiece for animal preparation and care and Scott Charlton for scanning assistance. This work was supported by the Canada Foundation for Innovation (to RSM); Canada First Research Excellence Fund to BrainsCAN; Brain Canada Platform Support Grant (to RSM); Natural Sciences and Engineering Research Council of Canada Discovery Grant (to RSM); and Canadian Institutes of Health Research FRN 148365 (to RSM) and FRN 353372 (to SE).

## Author Contributions

K.M.G., D.J.S., and S.E. conceived the social-coil method. K.M.G. and D.J.S. designed the restraint system. K.M.G. and A.M. fabricated the radiofrequency coil and acquired bench measurements. P.Z. designed and fabricated the custom platform. K.M.G., J.C.C., Y.H., J.S.G, and S.E. designed and conducted experiments. K.M.G. analyzed coil-performance data. J.C.C. and Y.H. analyzed functional data. K.M.G., J.C.C., Y.H., and D.J.S. wrote the manuscript. All authors reviewed the manuscript.

---

**Supplementary Fig. 1.**
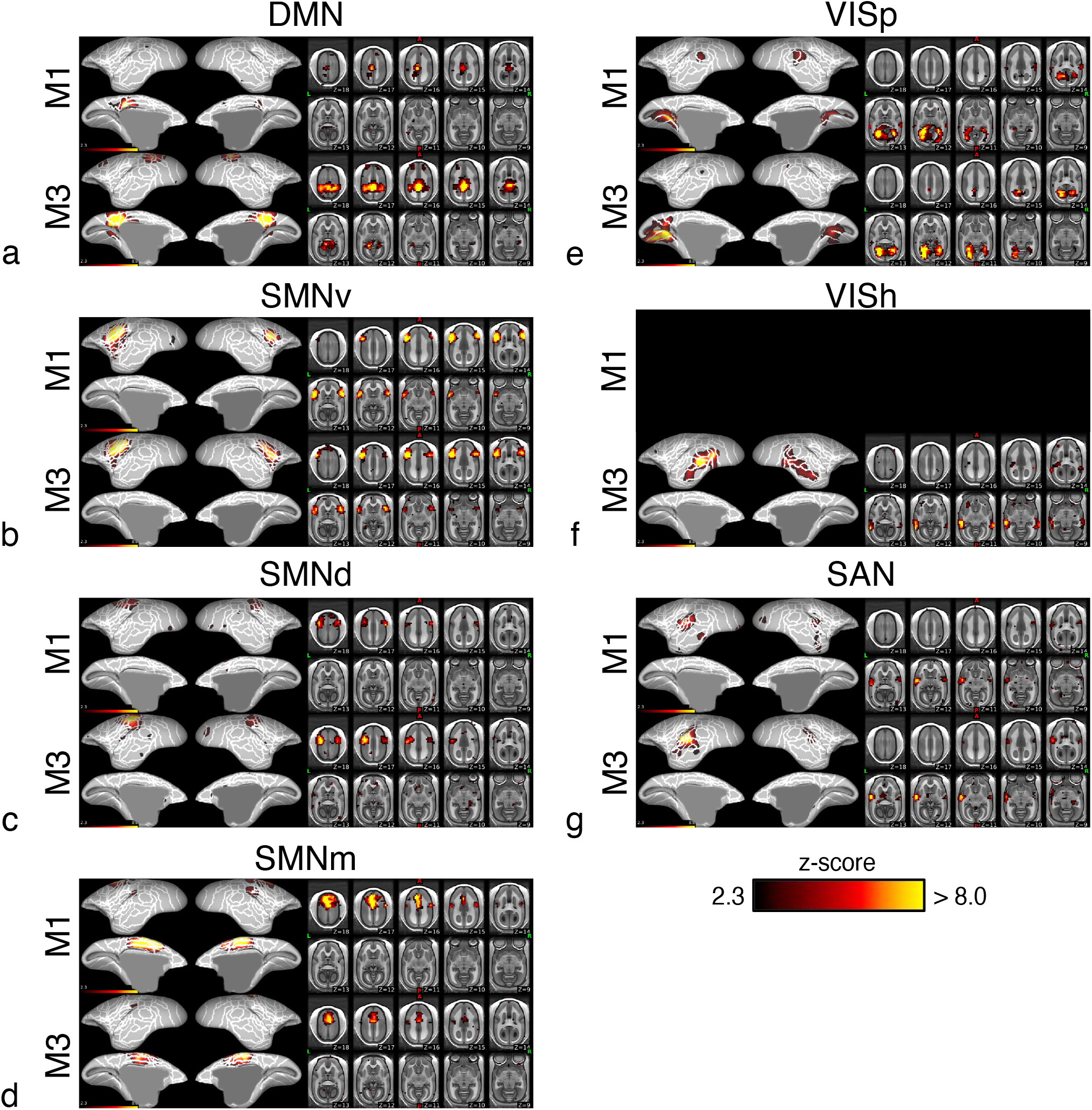
Functional networks present during constant interaction. Functional networks surpassing the significance threshold were: **a**, default mode network (DMN); **b**, ventral somatomotor network (SMNv); **c**, dorsal SMN (SMNd); **d**, medial SMN (SMNm); **e**, primary visual network (VISp); **f**, high-order VIS (VISh); and **g**, salience network (SAN). These networks are presented as *z*-score maps on the template surface and volume. Connectivity maps had similar distributions between the two marmosets, although VISh did not meet the significance threshold for monkey M1. White lines indicate cytoarchitectonic borders.

